# Overburdening of peer reviewers. A multi-disciplinary and multi-stakeholder perspective on causes, effects and potential policy implications

**DOI:** 10.1101/2021.01.14.426539

**Authors:** Anna Severin, Joanna Chataway

## Abstract

Peer review of manuscripts is labour-intensive and time-consuming. Individual reviewers often feel themselves overburdened with the amount of reviewing they are requested to do. Aiming to explore how stakeholder groups perceive reviewing burden and what they believe to be the causes of a potential overburdening of reviewers, we conducted focus groups with early-, mid-, and senior career scholars, editors, and publishers. By means of a thematic analysis, we aimed to identify the causes of overburdening of reviewers. First, we show that, across disciplines and roles, stakeholders believed that the reviewing workload has become so enormous that the academic community is no longer able to supply the reviewing resources necessary to address its demand for peer review. Second, the reviewing workload is distributed unequally across the academic community, thereby overwhelming small groups of scholars. Third, stakeholders believed the overburdening of reviewers to be caused by (i) an increase in manuscript submissions; (ii) insufficient editorial triage; (iii) a lack of reviewing instructions; (iv) difficulties in recruiting reviewers; (v) inefficiencies in manuscript handling and (vi) a lack of institutionalisation of peer review. These themes were assumed to mutually reinforce each other and to relate to an inadequate incentive structure in academia that favours publications over peer review. In order to alleviate reviewing burden, a holistic approach is required that addresses both the increased demand for and the insufficient supply of reviewing resources.

## 1. Introduction

Journal peer review is a process of scientific assessment by which manuscripts are evaluated by other scholars who are considered experts within the same or a related field (Severin and Chataway, 2020; Tennant et al., 2017). Peer review is expected to fulfil different functions, including conducting quality control, improving manuscripts, assessing the suitability of manuscripts, informing publication decisions, providing authors with feedback by their peers, curating academic communities and providing a seal of approval for publications (Severin and Chataway, 2020).

The traditional model of peer review has been recognized to put increasing strain on the academic community (Alberts et al., 2008). On average, 2.7 reviews are completed for every manuscript and writing a review takes a median five hours. An estimate of 13.7 million reviews are carried out per annum (Publons, 2018). It is widely assumed that the academic community is overburdened by this workload (Alberts et al., 2008; Arns, 2014; Stahel and Moore, 2014). Being overwhelmed with reviewing workload, reviewers might decline to review more often and editors might face difficulties in recruiting knowledgeable reviewers who agree to review, which could delay the review process. Also, reviewers might review in a haste and fail to detect errors in manuscripts (Elsevier and Sense about Science, 2019; Nicholas et al., 2015).

Innovations to peer review aimed at alleviating reviewing burden have been suggested and some are currently being tested. These include providing financial or non-financial incentives to reviewers, publishing review reports and revealing reviewer identities, training reviewers, artificial intelligence and machine learning aiding manuscript handling processes and reviewing, and databases of reviewers (Tennant et al., 2017; Van Noorden, 2014). The acceptance of these innovations within the academic community determines their success. Depending on how they perceive the reviewing burden and what they believe to be the underlying mechanisms that cause a potential overburdening of reviewers, stakeholders, including authors, reviewers, editors and publishers, might vary in their acceptance of potential solutions. This currently presents a knowledge gap as no research qualitatively explores stakeholders’ perceptions of reviewing burden. There are mathematical models that estimate the overall sustainability of the peer review system (Kovanis et al., 2016). Further, there are standardised surveys that gauge overburdening of reviewers as one of many potential reasons for why scholars decline to review (Brannon et al., 2016; Djupe, 2015; Mulligan et al., 2013; Publons, 2018; Tite and Schroter, 2007; Ware, 2008a, 2008b). Publishers and editors usually are not represented in these surveys and junior scholars are often under-represented. Qualitative analyses of stakeholder perceptions are another strand of research (Glonti et al., 2019; Harley et al., 2010; Severin and Chataway, 2020; University of Tennessee and CIBER Research Ltd, 2013). This research is not focused on reviewing burden, but indicates that stakeholders generally differ in their perceptions of challenges in peer review and respective solutions. For example, Zaharie and Osoian (2016) performed semi-structured interviews to explore the potential of incentives for improving peer review engagement. Senior scholars believed reviewing to be a reciprocal duty and hence did not expect to receive any rewards, while junior scholars perceived reviewing as a means of career advancement and appreciated being mentioned on the journal website (Zaharie and Osoian, 2016).

To our knowledge, no study comprehensively investigates how stakeholders perceive the burden that is placed on reviewers and the causes of a potential overburdening. This is important as considerable resources are dedicated to innovative review methods aimed at alleviating reviewing burden (Birgit and Edit, 2017). We conducted focus groups with stakeholders involved in academic publishing across academic disciplines to explore how junior- to senior-career scholars, reviewers, editors and publishers perceive reviewing burden (Severin and Chataway, 2020; Tennant et al., 2017).

## 2. The history and evolution of journal peer review

This chapter illustrates how peer review emerged in a response to the changing nature of academic research and communication.

### 2.1 The emergence and early history of peer review

Up until the 17^th^ century, feedback on academic work was predominantly sought through informal communication between authors and members of the academic community (Tennant et al., 2017). With the foundation of national academies in Europe, this was gradually replaced by a more formalised process. When academies created printed journals to curate research within their communities, they faced limited distribution capacities and had to choose work for publication. The process of independent review emerged, which was more a collegial exchange between authors and journal editors than a systematic mechanism by which external experts judged submitted work (Spier, 2002). What was accepted for publication remained a decision of the editors and those whose expertise they might or might not have consulted (Csiszar, 2016). This slowly changed when the Royal Society of London assumed editorial responsibility of *Philosophical Transactions* in the mid-18^th^ century. The society implemented a procedure by which manuscripts were scrutinized by a committee of society members knowledgeable in the field (Atkinson, 1999). Their assessment and recommendation to the editor were influential for the future of the manuscript. Other societies soon adopted similar practices for their journals, but journals without ties to societies were slow to follow (Spier, 2002).

### 2.2 Adaptation through professionalization and commercialization

In the 19^th^ century, the academic system experienced professionalization and specialisation. The numbers of scholars working in academia increased considerably, which led to a substantial increase in the quantity and specialization of manuscripts and a proliferation of scientific journals (Tennant et al., 2017). Facing these developments, editors began consulting expertise outside the immediate group of society committees. Peer review became a mostly outsourced process, which consulted external scholars knowledgeable in the topic of a manuscript (Csiszar, 2016). As the idea took hold that peer review ought to ensure the integrity of published manuscripts, independent journals too began establishing peer review (Spier, 2002).

In the second half of the 20^th^ century, editor-led peer review became mainstream. As the volume of manuscripts increased continuously, there was a need to be even more discriminative in selecting manuscripts for publication in printed journals and peer review gained importance as an objective judgement of research quality (Spier, 2002; Tennant et al., 2017). In addition, a prestige regime emerged around the perception of excellence of publishing in certain journals. These developments caused publishers to rely on peer review as a formalized process for determining which manuscripts are worth publishing (Tennant et al., 2017). These developments caused the demand for peer review to increase considerably. Based on the observation that it has become increasingly difficult to recruit reviewers, it was widely assumed that the increase in scientific production threatened the ability of the academic community to supply the reviewing resources necessary to address its own demand for peer review (Kovanis et al., 2016). Several voices raised concerns that, as a result, reviewers were overburdened (Arns, 2014; Brannon et al., 2016; Kovanis et al., 2016; Stahel and Moore, 2014).

The predominant peer review model in the 19^th^ and 20^th^ century was voluntary, anonymous and confidential. Voluntariness referred to the perception that peer review was a voluntary albeit reciprocal duty by scholars who devote their expertise and time without receiving financial compensation (Al-Khatib and Teixeira da Silva, 2019). Anonymity meant that the reviewer identity was hidden from the author (single blind review) or that both author and reviewer identities are unknown to each other (double blind review) (Ross-Hellauer et al., 2017). Confidentiality meant that reviewer identities and comments were not made public (Ross-Hellauer et al., 2017).

### 2.3 Diversification and innovation in modern peer review

Partly in response to a perceived overburdening of reviewers, several innovations to peer review have been suggested and some are currently being tested. One strand of innovation includes artificial intelligence and machine learning for increasing efficiencies in the manuscript handling process and for alleviating reviewing workload. This includes aiding manuscript handling, such as automatically identifying reviewers based on manuscript contents, assessing performance or conflicts of interest of reviewers and checking whether references and the manuscript structure meet journal policies (BioMed Central and Digital Science, 2017; Frontiers, 2018). It has also been suggested that automation processes could aid or replace review processes as such. This comprises plagiarism checks, identifying fraudulent behaviour and using language processing to extract key findings of a manuscript and placing these in context with existing work (BioMed Central and Digital Science, 2017; Frontiers, 2018; Heaven, 2018). A further strand of innovation is described as “open peer review”, which encompasses different ways in which peer review can be opened (Ross-Hellauer, 2017). First, this can include inviting the public to contribute or sharing review reports amongst reviewers to facilitate co-reviewing, thereby potentially distributing workload more equally and leveraging synergies. Second, this can include revealing author and reviewer identities and publishing review reports and author responses, in order to recognise reviews as scholarly outputs and publicly attributing these to their authors (Ford, 2013; Ross-Hellauer et al., 2017). Incentivising peer review to potentially motivate more scholars to agree to review describes another innovation that aims at alleviating reviewing burden. Incentives can be material in the form of reviewer payments, discounts on book purchases or fee waivers for publication in journals. Incentives can also be non-material, including giving credit for and displaying reviewing activities, either through external services or publishers (Ravindran, 2016).

## 3. Methods

By means of focus groups, we explored how stakeholders involved in peer review perceive current challenges in peer review and how these could be addressed, with a particular focus on the burden that is placed on reviewers. Methods are described below. A detailed account is published elsewhere (Severin and Chataway, 2020).

### Sampling

By means of maximum variation sampling (Breen, 2006), we selected participants who covered early- to senior scholars, reviewers, editors and publishers, and all academic disciplines, including social sciences, humanities, natural sciences and life sciences. We recruited participants from different sources, including university staff websites, journal editorial board websites, professional network websites and academic social networks. We contacted participants via email and provided them with a consent form and information sheet containing information about study aims, procedures, rules of conduct and confidentiality measures. We continued participant recruitment until saturation across sampling criteria was achieved (Severin and Chataway, 2020) (Table 1).

**Table 1:** Sampling criteria (Severin and Chataway, 2020)

| **Criterion** | **Description** |
| --- | --- |
| Professional background | Stakeholder involved in peer review processes at academic journals: Early-career scholars (PhDs, postdocs, lecturers, research fellows), senior scholars (assistant professors, professors and emeriti professors), editors and publishers |
| Journal characteristics | Scope (specialty journal and mega-journal); business model (open access, subscription-based and mixed), publisher (scholarly, commercial and mixed) |
| Academic discipline | Natural and life sciences, social sciences and humanities |
| Location | United Kingdom or Switzerland |

### Data collection

We held focus groups workshops in March to June 2019. Where possible, we structured workshops by stakeholder group to allow for similarity in participants’ experiences. Workshops were two-hour long and involved 3 to 7 participants each. Before each workshop, we restated study aims, procedures as well as confidentiality measures and obtained informed consent. We moderated discussions by means of a semi-structured topic guide, which was based on a review of the literature and further refined following a pilot workshop. We asked participants to discuss current challenges in peer review and how these could be addressed, with a particular focus on the burden that is placed on reviewers.

We audio-recorded all discussions, imported audio files to NVivo 12 and transcribed these. An assistant took notes (Severin and Chataway, 2020).

### Data analysis

We analysed transcripts thematically by exploring patterns and themes in relation to reviewing burden. Following an approach published elsewhere (Severin and Chataway, 2020), this was done in two steps. The first step included developing a preliminary codebook, which was driven by our research questions (AS and JC). In a second step, AS read and reread the transcripts and coded their topics. AS coded topics already entailed in the codebook while allowing new topics to emerge. AS repeated the coding until saturation across reviewing burden and potential causes was reached, defined as the point where no additional information was forthcoming from coding (Ando et al., 2014; Severin and Chataway, 2020). AS updated and revised the codebook continuously. Where codes emerged in a repeated pattern, they became a theme.

## 4. Results

A total of 37 participants were recruited for seven workshops (Table 2). This included five early-career researchers, four mid-career researchers, 17 senior researchers (of which 13 researchers also held an editorial position), eight publishers, and three editors who did not hold a position at a research institution. The groups of senior career scholars and editors are in large part overlapping as most recruited senior career scholars held an editorial position and because most editors held an academic position. Because both groups showed no differences in their perceptions, we will refer to them as one stakeholder group (Severin and Chataway, 2020).

**Table 2:** Participant characteristics

| Criterion | Description |
| --- | --- |
| Stakeholder group | Early-career scholars (n = 5), mid-career scholars (n = 4), senior-career scholars (n = 17), and editors (n = 3), publishers (n = 8) |
| Sex | Female (n = 12), male (n = 25) |
| Academic discipline | Natural and life sciences (n = 24), social sciences (n = 10), humanities (n = 6) and cross-disciplinary (n = 2) |
| Location | United Kingdom (n = 20), Switzerland (n = 17) |

When stakeholders were asked what they perceived to be important challenges in the peer review system, they agreed that the reviewing workload has become so enormous that the academic community might no longer be able to supply the reviewing resources necessary to address its own demand for peer review.

> “The volume of manuscripts that need to be reviewed is too much, too much to expect the top people in the field to read these thoroughly.” (Professor and editorial board member, mathematics)

Across disciplines and roles, stakeholders believed this imbalance between the demand for and the supply of reviewing resources to result in an overburdening of reviewers (Fig 1).

**Fig 1:**
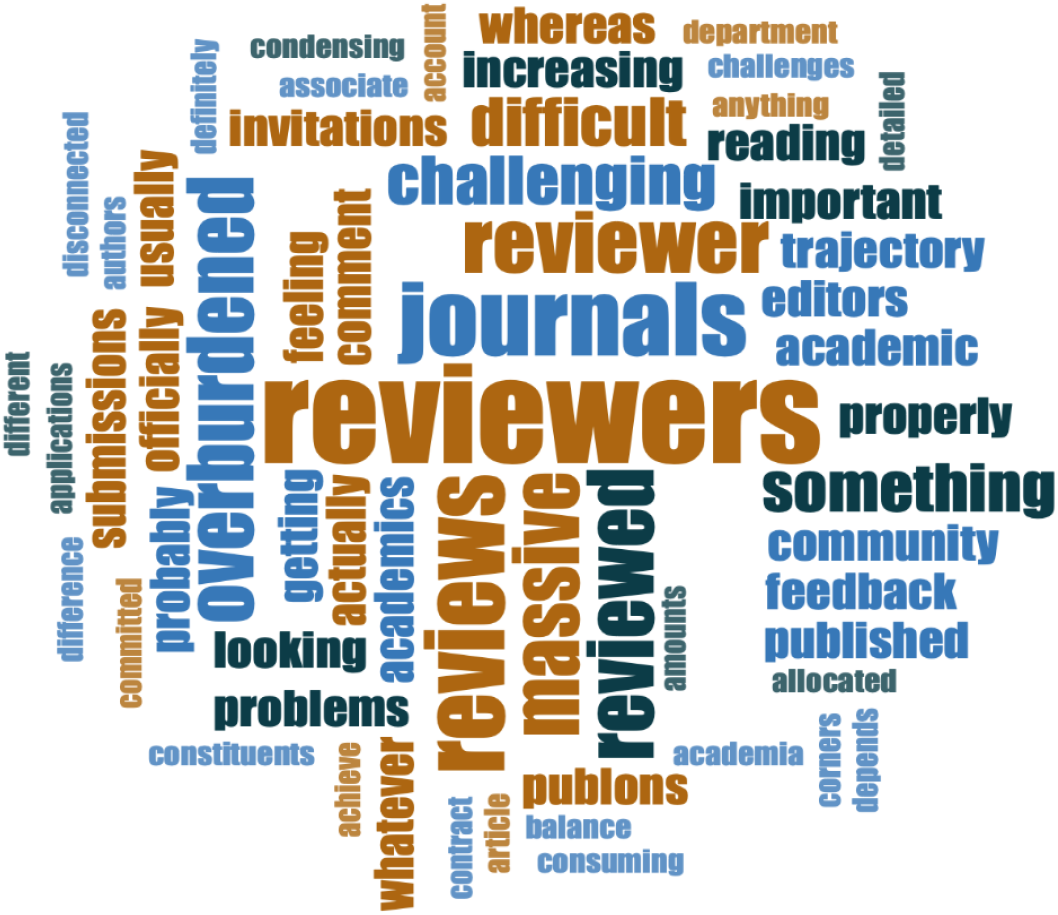
Word cloud of keywords. Keywords discussed in relation to overburdening of reviewers, represented by frequency (larger text indicates greater frequency) using NVivo 12.

**Fig 2:**
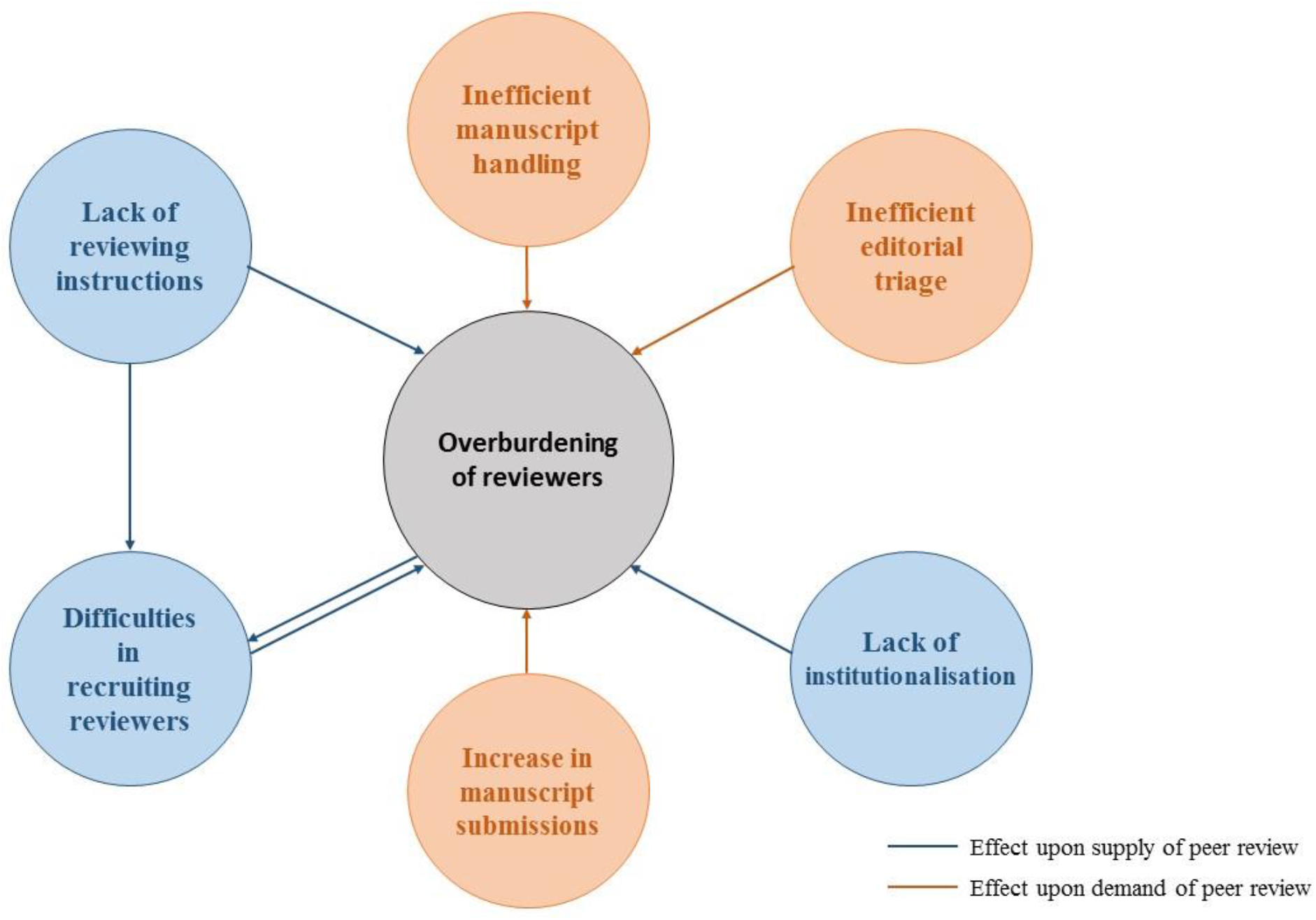
Causes of overburdening of reviewers

The overburdening of reviewers was perceived to be problematic for a number of reasons. To publishers and editors, the overburdening of reviewers became most visible in difficulties in recruiting reviewers for their journals.

> “As the volume of submissions that potentially will need to be reviewed is increasing, the time allocated to each individual reviewer is increasing. This strains the community. It is more challenging [to find reviewers].” (Publisher, cross-disciplinary)

Junior- to senior-career scholars and editors believed that reviewers occasionally might conduct superficial quality control. They assumed that, given their overwhelming workload, reviewers might not always have the time to read manuscripts thoroughly and might therefore fail to detect crucial errors in manuscripts.

> “The thing is that [scholars review] under huge time pressures because they have a full time job doing something else. They are squeezing this in to their full time job […] and therefore the temptation is that they will do a light quick easy job reviewing manuscripts. […] If they had more time, they would do a good job. I am sure they would, but very often, they do not have the time, so they will cut corners from time to time.” (Senior lecturer, computer sciences)

Delays in the review process were believed to be another consequence of the overburdening of reviewers. Junior- to senior scholars and editors saw the reason for this in reviewers not sticking to agreed timelines for submitting reviews.

> “My main point is that reviewers are on time. If they agree on a timeframe then they should comply with this timeframe. If this is three weeks or a month, or two months, if they comply with this timeframe, I am satisfied as an editor. Problematic are reviewers that are not corresponding to the time frame.” (Professor and editor-in-chief, political science)

Reflecting upon their own workload, mid- and senior scholars explained that delays in submitting reviews were natural as scholars might not always be able to balance heavy reviewing workloads with existing academic responsibilities. Junior scholars believed that editors also played a role in delaying the review process by failing to follow up on delayed reviewers.

### 4.1 Causes

Different themes were deduced in our analyses, which stakeholders believed to be causes of the overburdening of reviewers. These related to six key themes: (i) an increase in manuscript submissions; (ii) insufficient editorial triage; (iii) a lack of reviewing instructions; (iv) difficulties in recruiting reviewers; (v) inefficiencies in manuscript handling and (vi) a lack of institutionalisation of peer review. These themes were assumed to have an impact either upon the demand for or on the supply of reviewing resources. As shown in Fig. 1, these themes were also believed to be partly interrelated and to mutually reinforce each other.

### Increase in manuscript submissions

Across disciplines and roles, stakeholders believed that growing numbers of manuscripts submitted for publication were one of the main drivers of the overburdening of reviewers. The numbers of publications were perceived to have grown considerably across all disciplines, thereby increasing the demand for peer review, while the number of potential reviewers remained equal. Stakeholders concluded that the workload allocated to individual reviewers increased, causing reviewers to be overburdened.

> “Broadly speaking [there is] growth in the number of publications. […] so the volume of submissions that potentially will need to be reviewed is increasing and the time allocated to each individual reviewer is increasing” (Publisher, natural sciences)

While publishers did not speculate about the causes for the increase in manuscript submissions, scholars and editors presented a number of reasons they believed to be responsible. Across disciplines, these stakeholders felt that inadequate incentive structures were one of the main drivers for the increase in manuscript submissions. It was believed that, as the criteria for academic hiring and promotion as well as for research funding allocation increasingly focus on scientific publications, scholars are inclined to publish more. Particularly stakeholders from the social sciences and the humanities expressed concerns about the “publish-or-perish” culture of academia in which scholars have to publish work to advance their career. They added that, because reviewing remains unrecognised as a scholarly output in the academic career trajectory, scholars would further be discouraged from reviewing an adequate number of manuscripts from their peers in return for having their own work reviewed.

### Insufficient editorial triage

Across disciplines, stakeholders identified inadequate editorial triage as a further cause of the overburdening of reviewers. Given the increase of manuscript submissions, stakeholders expected editorial review to filter out manuscripts that are unlikely to survive the review process. By checking formality requirements, running plagiarism checks and evaluating if manuscripts meet minimum quality standards and the scope of a journal, the editorial team should decide whether manuscripts are forwarded into peer review or outright rejected. Stakeholders believed that too many unsuitable manuscripts are sent to review and concluded that editorial triage currently fails to regulate demand for peer review.

Reflecting on their roles as reviewers, junior- to senior-career scholars related this to the quality of manuscripts. They argued that peer review is overwhelmed with low-quality manuscripts.

> “Personally, I am surprised with the low quality of manuscripts that are sent to peer review. More papers should be outright rejected.” (Professor and editor, literature studies)

Scholars stated that this would mean avoidable work for them. Perceiving it an annoyance to review manuscripts of poor quality, some scholars stated that they started declining to review more often. Publishers, who expected editorial triage to assess the suitability of manuscripts for their particular journals, reported that too many papers are sent to peer review that are out of the scope of their journals.

> “I think it is a real challenge to try and match the editorial review with the aims and objectives of publication […]. So authors would be surprised in many cases when their article is accepted or rejected based on the stated aims of the journal […]. I think the big message is […] that it is problematic for publishers trying to get a handle on a focus for a particular community for their journal to meet the needs of that particular community.” (Publisher, cross-disciplinary)

Insufficient editorial triage was rationalised in different ways. Editors and senior scholars with editorial roles explained that, as their responsibilities were delegated to editorial teams and software systems, their own roles have been reduced considerably. While this allowed for a higher throughput of manuscripts, it has also reduced their oversight and their ability to filter out manuscripts that should not enter peer review. Reflecting upon their experience as authors, junior- and mid-career scholars explained that sometimes authors purposively submit manuscripts that are not yet publishable, hoping that they will be improved through peer review. As editors fail to filter out such manuscripts, reviewers would have to review these manuscripts, increasing their workload.

### Lack of institutionalisation

There was cross-disciplinary agreement amongst stakeholders that, because reviewing is not institutionalised, scholars face difficulties to balance reviewing with already existing responsibilities. Particularly scholars in the social sciences and humanities criticised that, even though reviewing serves crucial functions in scholarly publishing, it is neither part of their employment contract with the university nor included in research grants.

> “At the moment, what is weird is that there is a massive academic service that is really important for the whole community, but […] it is not officially part of my contract with the university, right? Technically, it is extra.” (Professor and editor, philosophy)

As a result, reviewing would rely on voluntary contributions by scholars who already have a full-time position that includes teaching and research responsibilities. Finding a way to engage in peer review on top of these duties poses a challenge.

> “[Scholars] review under huge time pressure because they’ve got a full time job doing something else, so they’re squeezing it in.” (Senior lecturer, computer sciences)

As scholars are requested to review growing numbers of manuscripts, they might not find the time to accommodate all requests to review and hence decline to review.

### Lack of reviewing instructions

Senior scholars with editorial positions, editors and publishers considered unclear reviewing instructions to be a further cause of the overburdening of reviewers. Stakeholders shared that reviewers might not always be well informed about the reviewing instructions of a journal. This was considered problematic because reviewers might not be able to correctly predict the amount of work it takes to review a manuscript. They might either underestimate the workload and be overburdened with the task, or overestimate it and decline the request to review, making it difficult for editors to recruit sufficient numbers of reviewers.

> “There is a perception around that reviewing is a very difficult task. […] I think that it would be better if the editors, or even on the journal website, stated what is required, as bottom line, of peer reviewers. […] I think if reviewers were aware of this it might be easier to get reviewers.” (Emeritus professor and editor-in-chief, mathematics)

It was also explained that in absence of clear reviewing guidelines, reviewers would assume what is expected of them. This might create redundant work as reviewers either assess aspects in detail that do not feed into publication decisions or fail to review aspects that are relevant, requiring editors to request additional comments.

> “I think this is central and I still have the same problem. You have to closely monitor the reasons for rejection. I sort of go back to them, “Are you sure about it really?” “(Publisher, engineering and technology)

Junior scholars agreed on the notion that there is a lack of clear reviewing instructions but did not relate this to the overburdening of reviewers.

### Difficulties in recruiting reviewers

Irrespective of their discipline, editors, mid- to senior-career scholars and publishers believed difficulties in recruiting reviewers to be a driver of the overburdening of reviewers. They believed that, as only certain groups of scholars engage in peer review, reviewing workload would be distributed unequally across members of the academic community. The reviewing burden carried by some groups of scholars might be so high that it is potentially unmanageable, likely causing them to be overwhelmed and overworked.

It was reported that authors located in Asian countries submit growing numbers of manuscripts but are less often invited to review than reviewers located in high-income countries, particularly North America and Europe. Further, because many journals consider having published previously as a requirement for being qualified as a reviewer, editors might not recruit early-career scholars, even though they would be willing and capable to provide thorough reviews. It was believed that, as editors and publishers currently fail to recruit authors located in Asian countries and early career scholars as reviewers, the reviewing workload is skewed geographically and demographically.

Stakeholders assumed these inequalities to be intensified by a lack of clear reviewing instructions. Particularly editors believed that, because reviewing criteria are not always clear to potential reviewers, inexperienced scholars might overestimate the effort involved in reviewing and decline to review. Senior scholars and editors added that inadequate incentive structures also exacerbate inequalities in the distribution of reviewing burden. It was argued that, because the criteria for academic hiring and promotion as well as for funding allocation put most weight on publications, scholars who still have to secure tenure would be inclined to publish their own research without reviewing an adequate number of their peers’ manuscripts.

> “Of course, reviewing goes into Research Excellence Framework statement […], but it is sort of weirdly disconnected […] I think it would matter if there were a way that reviewing would be taken into account in all the actual things that matter to the academic trajectory, right? So, in the context of funding, the career and the tenure, and all these things. Because that is the thing that really matters to academics” (Professor and editor, Philosophy)

Generally, stakeholders assumed that difficulties in recruiting reviewers and the overburdening of reviewers would mutually reinforce each other. Stakeholders argued that, as reviewers face an overwhelming workload, they would decline to review more often, making it difficult for editors to recruit sufficient numbers of reviewers.

> “The volume of submissions that potentially need to be reviewed is increasing the time allocated to each individual reviewer and you are looking at the difference between two invitations being sent to three invitations being sent to get one review back over a three year period between 2013 and 2016.” (Publisher, cross-disciplinary)

As a result, those groups of scholars who are still willing to engage in peer review would face an ever-increasing workload, adding to the risk of being overwhelmed.

Junior scholars did not perceive reviewer recruitment as an important driver of reviewer overburdening but shared their experiences with reviewers not being available for all rounds of revision. Often, editors would then recruit new reviewers who might request additional revisions, causing the publication of their manuscripts to be delayed.

### Inefficient manuscript processing

Taking a holistic approach, publishers, editors and senior scholars with editorial positions stated that inefficiencies in the overall manuscript handling process create redundant work for reviewers.

One inefficiency that stakeholders identified was that peer review is not re-used when a manuscript has been rejected by one journal and is then submitted to another journal. Usually, when a rejected manuscript is submitted to elsewhere, editors recruit reviewers to assess the manuscript again and new demand for peer review is created, irrespective of the fact that a manuscript might have been thoroughly assessed before.

> “The same journals ask the same people to review a paper that has been rejected in one place and then goes to somewhere else. […] Sometimes, if the paper has been rejected in one journal, the peer review is still really useful […]. And that peer review does not get carried forward. There is a lot of inefficiencies.” (Publisher, cross-disciplinary)

Stakeholders further pointed out that basic manuscript processing steps are still performed manually, even though they could be automated. Examples mentioned by stakeholders included identifying reviewers and correspondence with reviewers and authors, assessing conflicts of interest of reviewers and checking the manuscript structure meets journal policies. These inefficiencies were believed to create additional albeit avoidable work for reviewers and editors, thereby further overwhelming the academic community.

## 5. Discussion and conclusion

By means of qualitative focus group discussions, this study provided an in-depth exploration of how stakeholders involved in peer review, including early-, mid-, and senior career scholars, reviewers, editors, and publishers, perceived the burden that is placed on reviewers and what they believed to be the drivers of a potential overburdening of reviewers. It was also important to examine whether stakeholder perceptions differed depending on their disciplinary background or their relationship with the process.

One key finding of this study was that across roles and disciplines stakeholders believed the reviewing workload to have become so enormous that it has reached a breaking point beyond which the academic community is no longer able to supply the reviewing resources necessary to address its own demand for peer review. To stakeholders, this showed in reviewer fatigue, that is, individual reviewers being overburdened with the amount of reviewing they are requested to do. This finding is in line with anecdotal evidence of overburdened and overworked reviewers (Alberts et al., 2008; Arns, 2014; Stahel and Moore, 2014). It also confirms reports showing that reviewers increasingly decline to review, which limits publishers and editors in their ability to recruit sufficient numbers of qualified reviewers (Brannon et al., 2016; Breuning et al., 2015; Djupe, 2015; Ellwanger and Chies, 2020; Fox et al., 2017; Mulligan et al., 2013; Publons, 2018; Tite and Schroter, 2007; Ware, 2008a). This study revealed a more differentiated account of the distribution of reviewing burden than most anecdotal reports of reviewing burden currently do. Stakeholders perceived that small groups of scholars would carry a disproportionate part of the overall reviewing workload, which might result in individual reviewers being overwhelmed and potentially overworked. This finding confirms mathematical models of the overall sustainability of the peer review system, which revealed that a small number of researchers handles a large share of the overall reviewing workload (Kovanis et al., 2016; Publons, 2018).

As a further key finding, this study showed that the overburdening of reviewers is caused by an imbalance between the demand for and the supply of reviewing resources. The underlying causes of this imbalance were related to the incentive structure of academia. Stakeholders believed that, as the criteria for academic hiring and promotion as well as for the allocation of research funding prioritise publications over peer review, scholars might be inclined to publish their own research without reviewing an adequate number of their peers’ manuscripts in return. Particularly stakeholders from the social sciences and the humanities expressed concerns about the consequences of the “publish-or-perish” culture of academia in which scholars have to publish academic work to further or maintain their academic career. This means that in order to alleviate reviewing burden, it is necessary to change the overall incentive structure in academia. This could be done by recognising peer review as a scholarly output and letting it inform academic hiring and promotion decisions as well as the allocation of research funding.

A further key finding was that the causes of reviewer overburdening are interrelated and, in part, mutually reinforce each other. Such complex interdependencies again stress the need for adopting a holistic approach in alleviating reviewing burden.

Finally, depending upon their experiences and their relationship with the review process, stakeholders put different weight on the themes mentioned above as causes of overburdening, but did not disagree in principle. This means that it might be challenging but not impossible to find solutions that are acceptable to the wider community.

We recognize a number of limitations in our study. First, there are limitations related to our sampling approach. Because participants were recruited using purposive maximum variation sampling, there might have been selection biases in how we selected participants. We tried to alleviate this by means of pre-defined recruitment criteria. Further, because focus groups were face to face, stakeholders who lived far from the workshop location were less likely to join than stakeholders within close proximity were. We compensated travel costs to reduce geographical biases. Nonetheless, stakeholders based in other countries than Switzerland or the England were not represented in this study. Because academic publishing differs geographically (Collyer, 2018; Severin and Chataway, 2020), the generalizability of results was limited. Moreover, due to limited resources, the size of our sample was comparatively small, which might have limited our capability to comprehensively depict the causes of reviewer overburdening from the viewpoints of all relevant stakeholders. Second, because the validity of self-reported attitudes might suffer from inaccuracies in recollection, erroneous perceptions, incapability to answer correctly, and socially desirable answering (O’Sullivan, 2008), there might be inconsistencies between what stakeholders stated to be the drivers of overwhelmed reviewers and what they actually believed to cause overburdening. Finally, qualitative studies always includes some degree of subjectivity as the researcher’s experience and judgement influence how data are collected, analysed and interpreted. To mitigate subjectivity, we based our analysis and interpretation on a codebook and give exemplary participant quotes (Severin and Chataway, 2020).

To our knowledge, this study is the first to explore how different stakeholder groups experience the reviewing burden that is placed on scholars and where they identify the causes for a perceived overburdening. Having identified underlying mechanisms of the overburdening of reviewers, this study aids understanding reviewing burden as an important challenge in the current peer review system. Based on this understanding, potential solutions can be developed and implemented.

## Notes

### Competing Interest Statement

The authors have declared no competing interest.

